# Programmed Cell Death in the Developing *Brachypodium distachyon* Grain

**DOI:** 10.1101/775833

**Authors:** Safia Saada, Charles Ugochukwu Solomon, Sinéad Drea

**Author notes:** Author for correspondence; Charles Ugochukwu Solomon.

## Abstract

- The normal developmental sequence in a grass grain entails the death of several maternal and filial tissues in a genetically regulated process termed programmed cell death (PCD). The progression and molecular aspects of PCD in developing grain have been reported for domesticated species like barley, rice, maize and wheat. Here, we report a detailed investigation of PCD in the developing grain of a wild model species, *Brachypodium distachyon*.
- We detected PCD in developing *Brachypodium* grains using molecular and histological approaches. We also identified and surveyed the expression of *Brachypodium* orthologs of protease genes known to contribute to grain PCD.
- We found that *Brachypodium* nucellus degenerates by PCD in a centrifugal pattern following anthesis, although at a slower rate compared to cultivated cereals. Mesocarp PCD was not coordinated with endosperm development. *Brachypodium* lacks an expansion of vacuolar processing enzymes known for their roles in nucellar PCD.
- Combined with existing knowledge on grain PCD, our study suggests the importance of rapid nucellar PCD for grain size and that the pattern of mesocarp PCD affects grain shape.

## Introduction

Ordered and finely regulated elimination of cells in living organisms is called programmed cell death (PCD). PCD is a hallmark of numerous plant developmental processes including cereal endosperm development (Daneva *et al*., 2016; Domínguez & Cejudo, 2014; Kacprzyk *et al*., 2011). At fertilization, the fusion of a sperm nucleus with two polar nuclei gives rise to a triploid endosperm. At this point the nascent cereal endosperm is embedded within the nucellus, enclosed by two integuments, surrounded by the pericarp. However, by the time the grain is mature, these maternal tissues, as well as starchy endosperm cells have undergone PCD.

The post-fertilization series of PCD events during cereal grain development is well reported in barley, maize, rice and wheat. Generally, the nucellus rapidly undergoes PCD within the first five days after fertilization (Domínguez & Cejudo, 2014). PCD-degenerated nucellus cells provide growth space and resources for the incipient endosperm at this stage (Domínguez & Cejudo, 2014; Morrison *et al*., 1978; Norstog, 1974). In maize, nucellus PCD proceeds acropetally while in barley, rice and wheat, nucellus PCD is centrifugal (Chen *et al*., 2015; Radchuk *et al*., 2010; Yin & Xue, 2012). All that remains of the nucellar tissue at about 6 days post anthesis (DPA) are; 1. nucellar epidermis; bordered on the outside by the inner integument, 2. nucellar lysate; debris of lysed nucellus cells, which is sandwiched between the nucellar epidermis and the expanding endosperm, and 3. nucellar projection; located at the chalazal region which acts as transport route for nutrient supply to the endosperm (Evers & Reed, 1988; Freeman & Palmer, 1984; Morrison *et al*., 1978; Norstog, 1974; Oparka & Gates, 1981; Wang *et al*., 1994). In barley and wheat the nucellar epidermis persists for a longer period during grain development, retaining cytological integrity and metabolic activity until about halfway through grain development (Domínguez & Cejudo, 1998; Norstog, 1974). In rice the nucellar epidermis serves as transport route for nutrients into the endosperm and eventually collapses by 21 DPA, terminating grain filling (Ellis & Chaffey., 1987; Oparka & Gates, 1981; Wu *et al*., 2016). Although the nucellar epidermis is dead and crushed between aleurone and inner integument in mature temperate cereals (Hands & Drea, 2012), the time point of its PCD during grain development has not been reported.

Barley, rice and wheat nucellar projection are functional equivalents of placento-chalazal layer of maize and sorghum. Both tissues are located between the main vascular bundle and the developing endosperm. They differentiate into transfer cells that conduct nutrients towards the endosperm (Domínguez & Cejudo., 2014). PCD in nucellar projection/placento-chalazal region proceeds from cells proximal to the endosperm towards cells at the chalazal region (Kladnik *et al*., 2004; Thiel *et al*., 2008; Wang *et al*., 1994). PCD was detected in rice nucellar projection at about 3 DPA, evident in barley, maize and sorghum nucellar projection/placento-chalazal cells at about 8 DPA, and was detected in wheat at 13 DPA (Dominguez *et al*., 2001; Kladnik *et al*., 2004; Radchuk *et al*., 2010; Thiel *et al*., 2008; Yin & Xue, 2012). The degeneration of nucellar projection/placento-chalazal region creates transient sinks called endosperm cavity in barley and wheat, and placental sac in sorghum and some varieties of maize (Wang *et al*., 1994).

At anthesis, the cereal pericarp, from outside to inside, is composed of a layer of cutinous epicarp, several layers of parenchymal mesocarp and 2-3 layers of chlorenchymal endocarp. PCD occurred in barley and rice mesocarp cells closer to the integuments at 6 DPA and proceeded in an outward pattern. By 15 DPA, the mesocarp and endocarp have undergone PCD leaving behind two to three layers of empty cells that are crushed between the outer cuticle and the integument (Domínguez & Cejudo, 2014; Freeman & Palmer, 1984; Wu *et al*., 2016). While PCD in maternal nucellar and pericarp tissues of cereals generally results in post mortem disintegration, and remobilization of the cellular remains by filial endosperm and embryo, PCD leaves the starchy endosperm cells intact (albeit dead) until germination, when they are degraded and their contents used to fuel the growth of the embryo. Endosperm PCD was detected at about 8 DPA in rice, and 16 DPA in maize and wheat. While the progression of endosperm PCD is centrifugal in rice, basipetal in maize, wheat endosperm PCD proceeds randomly (Young *et al*., 1997; Kobayashi *et al*., 2013; Wu *et al*., 2016). Progress has been made in elucidating molecular mechanisms underlying PCD in developing cereal grains. A number of proteolytic genes belonging to different protease families have been linked with PCD of specific tissues in developing cereal grains. The expression profile of nucellin, an aspartic protease that belongs to Clan AA Family A1 (http://merops.sanger.ac.uk), highly correlates with the degeneration of the nucellus in barley and rice, prompting the conclusion that it has a role in nucellus PCD (Bi *et al*., 2005; F. Chen & Foolad, 1997). Another known aspartic protease Oryzasin (Os05g0567100), was found to be expressed during rice seed ripening and germination (Asakura *et al*., 1995). Notably, cysteine peptidases belonging to the legumain family (C13), also known as vacuolar processing enzymes (VPEs) have been shown to be expressed in timely and tissue specific manner, that coincides with caspase-like activities in developing barley grains (Borén *et al*., 2006; Julián *et al*., 2013; Linnestad *et al*., 1998; Radchuk *et al*., 2010; Sreenivasulu *et al*., 2006; Tran *et al*., 2014). The mRNA of barley nucellain orthologs was shown to localize *in situ* on nucellar cells undergoing PCD in wheat and *Brachypodium distachyon* (Drea *et al*., 2005; Opanowicz *et al*., 2011). Rice nucellain, *OsVPE1* was significantly down-regulated in OsMADS29-knock-down transgenic seeds which may indicate direct regulation of *OsVPE1* by *OsMADS29* in rice (Yang *et al*., 2012). Recent functional characterization of *HvVPE4* by RNA interference revealed that pericarp PCD is inhibited by downregulation of *HvVPE4* leading to reduced size and storage capacity in the embryo and endosperm (Radchuk *et al*., 2018). The expression of a cathepsin B-like protease gene belonging to the papain family (C1), overlaps with PCD in nucellus, nucellar epidermis and nucellar projection of developing wheat grains (Domínguez & Cejudo, 1998). A rice Cys protease gene, Os02g48450, also belonging to the papain (C1) family is severely downregulated in the nucellar projection of *A-OsMADS29* (*OsMADS29-knockdown*) lines. Promoter analysis of Os02g48450 upstream sequence revealed clusters of CArG-box motifs that was experimentally shown to be bound by *OsMADS29* (Yin & Xue, 2012). The rice MADS box family transcription factor *OsMADS29*, an ortholog of *Arabidopsis thaliana TRANSPARENT TESTA 16* (*TT16*), regulates PCD in the pericarp, ovular vascular trace, integuments, nucellar epidermis and nucellar projection, during rice grain development. Suppression of *OsMADS29* expression either by antisense constructs or RNA interference resulted partially filled grains that were small in size and shrunken in shape. The knockdown mutant grains also had smaller endosperm cells, reduced starch synthesis and abnormally shaped starch granules (Nayar *et al*., 2013; Yang *et al*., 2012; Yin & Xue, 2012).

*Brachypodium distachyon* (subsequently *Brachypodium*) recently became established as a model system for grasses. It is phylogenetically related to wheat, barley and rye. Mature *Brachypdoium* grains have less starch in thick walled endosperm cells compared to cereal grains. This is unusual because *Brachypodium* expresses the full set of genes required for starch synthesis in its endosperm (Trafford *et al*., 2013). Other aspects of grain development in B. distachyon have been reported, furnishing evidence for the suitability of *Brachypodium* grains as a system for comparative grain evolution and development relevant to tropical and temperate cereals (Francin-Allami *et al*., 2019; Francin-Allami *et al*., 2016; Guillon *et al*., 2011; Hands *et al*., 2012; Hands & Drea, 2012; Kourmpetli & Drea, 2014; Opanowicz *et al*., 2011; Trafford *et al*., 2013). However, details of PCD events, patterns and progression in developing *Brachypodium* grains to the best of our knowledge has not yet been reported.

From a previous study (Hands *et al*., 2012), we learned that *Brachypodium* grains have enlarged persistent nucellar epidermis at maturity. This contrasts other cereal grains where degeneration of nucellar epidermis and other maternal tissues leaves room and provision for the expanding endosperm. We therefore hypothesized that the pattern and progression of PCD in *Brachypodium* grains may be different from cereals. Here, we undertook a systematic histochemical and molecular study of PCD in developing *Brachypodium* grains. In addition, we surveyed and validated the expression profiles of B. distachyon genes belonging to protease families; A1, C1 (papain), C13 (VPEs) and C14 (Metacaspases) in an RNA-Seq dataset of developing *Brachypodium* grains. Protease genes belonging to these selected families have been linked to PCD events in developing grains of other species as reviewed above.

In this study, our results indicate that the rate of nucellar PCD is slow in *Brachypodium* compared to cereals. On the other hand, post mortem clearance of the mesocarp cells proceeds more rapidly in *Brachypodium* compared to cereals. Gene expression analysis suggests conserved roles of *Brachypodium* orthologs of proteases previously known to be involved in PCD and also yielded new candidate genes that may be part of the *Brachypodium* grain PCD molecular machinery.

## Materials and Methods

### Plant materials

*Brachypodium* grains (Bd 21), were imbibed on moist filter paper in a Petri dish and left at 5°C for two days to vernalize. They were transferred to room temperature (about 25-27°C) and left to germinate. After 7 days, the most virile seedlings were transferred to 9:1 Levington M2 Pot and Bedding Compost: Levington Fine Vermiculite mix (http://dejex.co.uk), in Vacapot 15 on plastic seed trays (www.plantcell.co.uk). They were grown in the greenhouse at 16hr daylight and 25°C temperature. The plants were regularly watered manually. Pre-anthesis ovary samples were collected at yellow (intact) anther stage. Grain samples were collected from spikes staged at anthesis.

### TUNEL staining

DNA fragmentation was detected using Terminal deoxynucleotidyl transferase (TdT) dUTP Nick-End Labeling (TUNEL). The assay was performed on dewaxed and rehydrated sample slides treated with proteinase K, according to the manufacture’s protocol of the *in situ* Cell Death Detection kit (Roche Diagnostics, Germany). Imaging was done with Nikon ECLIPSE 80i fluorescence microscope (Nikon, Japan), having an LED-based excitation source (CoolLED, presicExcite), using Nikon Plan Fluor 10x/0.30 DIC L/N1 objective lens. Fluorescence images were captured with a DS-QiMc cooled CCD camera (Nikon, Japan). Images were previewed, captured and saved using NIS-Elements Basic Research v3.0 software (Nikon, Japan) in JPEG2000 format.

### DNA isolation and electrophoresis

Genomic DNA was isolated from grains according to CTAB protocol. DNA extracts were quantified with nanoDrop spectrophotometer (Thermo Fisher Scientific). A 15 μl aliquot of 300 ng/μl of DNA was separated on 1% agarose gel at 100 V for 1.50 hr.

### Evans Blue staining

Freehand sections (10 – 30 μm) of freshly sampled *Brachypodium* grains were immersed in 2 ml Evans Blue solution (0.05% in water) for 3 min and rinsed in several changes of distilled water with gentle agitation for mortal staining. Stained sections were mounted in 50% glycerol, viewed and photographed with Zeiss Stereo Microscope equipped with a GT-vision GXCAM-5MP digital USB camera and GXCAPTURE software.

### Thin section and light microscopy

About 1 mm middle grain transverse sections were obtained from 5 DPA grains of *Brachypodium* and barley (cv. Bowman) (grown in same conditions as *Brachypodium*), under a dissecting microscope. Sections were fixed in 2.5% glutaraldehyde in 0.1M sodium cacodylate buffer pH 7.4, for 3 days at 4°C with constant gentle agitation and then washed in 0.1 M sodium cacodylate buffer. Further fixation in 1% aqueous osmium tetroxide was followed by dehydration in series of increasing Ethanol concentrations followed by propylene oxide. The sections were embedded in Spurr’s hard resin and polymerised for 16 hours at 60°C. 400 nm thick sections were obtained with ultramicrotome, stained for 30s with 0.01% toluidine blue, mounted in resin, and imaged with GX L3200B compound microscope (http://gtvision.co.uk) equipped with a CMEX-5000 USB2 Microscope camera and ImageFocus 4 software (http://euromex.com).

### Source and phylogenetic analysis of selected protease family genes

*Brachypodium* protein sequences belonging to *Brachypodium* protease families A1, C1, C13 and C14 were retrieved using INTERPRO (Mitchell *et al*., 2018) and PANTHER (Thomas *et al*., 2003) accession number specific for each domain (A1: pepsin; IPR001461, PTHR13683), (C1: papain; IPR000668, PTHR12411), (C13: legumain; IPR001096, PTHR12000), (C14: caspase; PTHR31773, PTHR31810) from the peptidase database MEROPS 12.1 (Rawlings *et al*., 2011). These accession numbers were also used to blast Phytozome v.12 (https://www.phytozome.jgi.doe.gov) and EnsemblePlants (http://plants.ensembl.org). The protein sequences were uploaded into Geneious R10 (https://www.geneious.com) and filtered by selecting only the unique longest version of the protein sequences. The same approach was used to obtain A1 and C1 protein sequences of *Arabidopsis*, barley, rice and wheat. Protein sequences of C13 and C14 of *Arabidopsis*, barley and rice were gathered from published reports on these proteins for each species (Bostancioglu *et al*., 2018; Julián *et al*., 2013; Rocha *et al*., 2013; Vercammen *et al*., 2004; Wang *et al*., 2018). Further information on sequences used in phylogenetic analysis is provided in the Table S2. Phylogenetic analysis was performed in MEGA6 (Tamura *et al*., 2013) and Geneious R10. Alignment was performed using MAFFT v7.308 (Katoh & Standley, 2013) under the following parameters: BLOSUM62 as scoring matrix, Gap open penalty of 1.53 and an offset value of 0.123. Parameters used to construct A1 and C1 phylogenetic trees are provided in the appendix. Phylogenetic tree of C13 was constructed using Maximum Likelihood method based on the Le_Gascuel_2008 model (Le & Gascuel, 2008). Phylogenetic tree of C14 was constructed using Maximum Likelihood method based on the Whelan and Goldman model (Whelan & Goldman, 2001).

### RNA-seq data source, processing and expression analysis of protease genes

RNA-Seq data of developing *Brachypodium* grain generated in our lab and publicly available at E-MTAB-7607 (http://ebi.ac.uk/arrayexpress) was used. Data quality was checked with FastQC. Reads were mapped with STAR v2.5.2b (Dobin *et al*., 2013), to *Brachypodium distachyon* v3.0.dna.toplevel.fa downloaded from Phytozome v.12. Mapped reads were counted with featureCount function from Rsubread v 1.22.2 package (Liao *et al*., 2019). Raw read counts were normalized with DEseq2 v1.6.3 (Love *et al*., 2014). Reads were scaled using unit variance scaling method. The expression profile of protease gene families in the RNA-Seq data was visualized with the package ComplexHeatmap v1.99.8 (Gu *et al*., 2016).

### mRNA *In situ* Hybridisation

Samples from five stages of grain development were harvested, processed and prepared for mRNA in situ hybridization as described previously (Drea, Corsar *et al*., 2005; Opanowicz *et al*., 2010). The probe template consisted of *BdMADS29* cDNA fragment amplified with gene specific primers (see Table S3 for primer sequences) and transcribed *in vitro* with T7 RNA polymerase.

### Real-time quantitative PCR

Primer sequences are provided in Table S2. Quantitative RT-PCR reactions were carried out using SYBR Green (SensiMix SYBR Low-ROX Kit) (BIOLINE) on 7500 MicroAmp Fast Optical 96-Well Reaction Plate (Applied Biosystems), with the following reaction mix components (5 μl of 2x SensiMix SYBR Low-ROX, 0.1 μl of 25 μM Forward/reverse Primers, 3.8 μl of DNase-free H_2_0, 1 μl of cDNA template). The thermal cycling conditions started by polymerase activation at 95°C for 10 min followed by 40 cycles of 95°C for 15 seconds then the annealing temperature of 65°C for 15 seconds flowed by 72°C extension for 15 seconds. Samples were normalized using BdACT7 expression and relative expression levels were determined using the 2^(-ΔΔCt) analysis method described in (Livak & Schmittgen, 2001) with 7500 Software v2.0.6 Applied Biosystems.

## Results

### TUNEL and vital staining reveal pattern and progression of PCD in developing *Brachypodium* grain

One feature of PCD is the disintegration of genomic DNA to fragments less than 200 nt. The 3’ OH end of these fragments can be labelled in situ by the TUNEL assay. These fragments can also be detected as laddering patterns when DNA from tissues undergoing PCD is analysed by gel electrophoresis. Evans blue is a vital stain that is excluded from living cells by intact cellular membrane but stains dead and dying cells that have lost membrane integrity. Evans Blue have previously been used to study PCD in cereals (Young & Gallie, 1999; Young *et al*., 1997). TUNEL-positive signals and Evans blue staining of sections made from unfertilized *Brachypodium* ovary (pre-anthesis ovary) (Fig. 1a,b; Fig. S1) suggests that PCD of nucellar and mesocarp cells precedes fertilization. Nucellus PCD progressed in a centrifugal pattern post-fertilization. Although, TUNEL-positive signal and Evans blue staining suggest rapid PCD of the nucellus, post-mortem clearance of dead nucellus cells seems to progress slowly. Whereas we observed complete clearance of nucellus cells at 5 DPA in barley, four to six layers of nucellus cells is still present in 5 DPA *Brachypodium* grain (Fig. 2). The nucellar epidermis of *Brachypodium* grains serves as an assimilate transport route during grain filling and is greatly enlarged by 5 DPA (Fig. 2a). TUNEL and Evans blue staining indicated nucellar epidermis PCD at 10 DPA (Fig. 1g,h; Fig. S1d). Integument PCD was detected as early as 2 DPA appears to continue up to 6 DPA (Fig. 1c-f).

**Figure 1.**
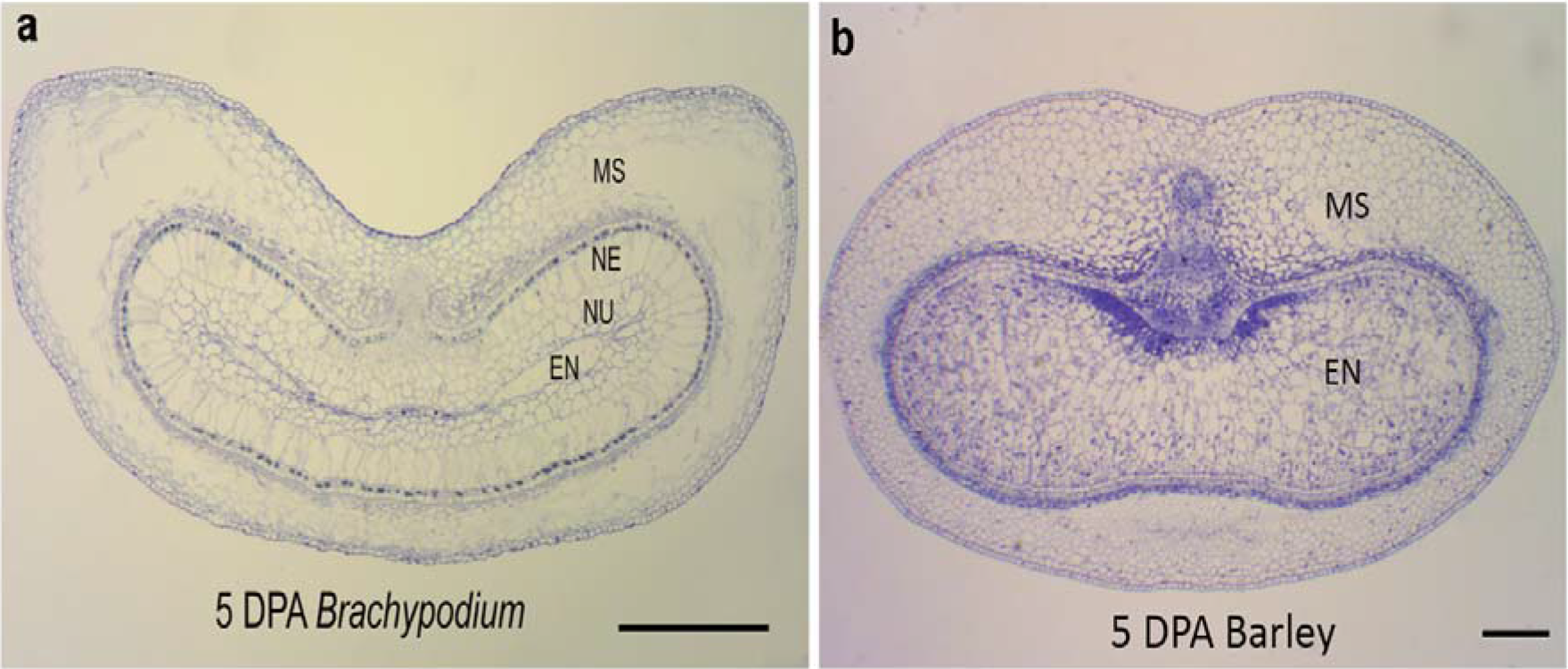
DNA fragmentation detected in *Brachypodium* grain section in five developmental stages,(a) Pr As Ov: pre-anthesis ovary, (c) young grain at 2 days post anthesis (2 DPA), (e) mid-length grain at 6 DPA, (g) full-length grain at 10 DPA, (i) mature grain at 20 DPA, (b-d-f-h-j) negative control, NE: nucellar epidermis, Nu: nucellus paremchyma cells, Mc: mesocarp, Pa: palea, In: integument, En: endosperm, Scale bar 100 μm.

**Figure 2.**
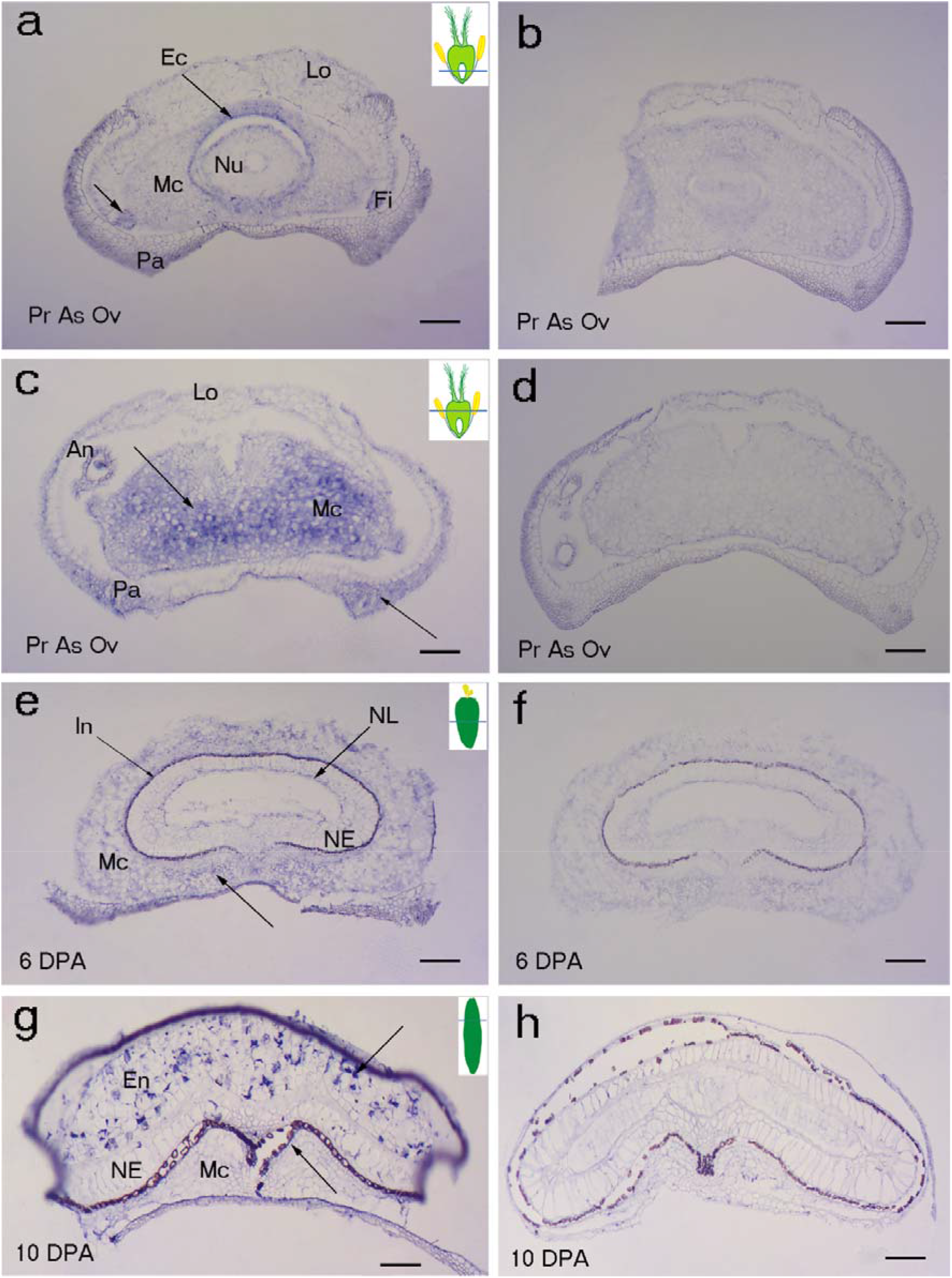
Comparison of grain development in *Brachypodium* and barley. At 5 DPA, (a) *Brachypodium* endosperm is barely differentiated and is surrounded by several layers of nucellar cells. Also, mesocarp cells have degraded by PCD. (b) Barley endosperm is fully differentiated and set for filling. Nucellar cells are absent and mesocarp cells appear intact. Scale bars: 0.2 mm. EN, endosperm; MS, mesocarp; NE, nucellar epidermis, NU, nucellus.

Differentiation of transfer cells (pigment strand and nucellar projection) that channel assimilates from the grain vascular bundle to the endosperm is evident in young grain sections because they exclude Evans Blue stain (Fig. S1b). The transfer cells are not stained by Evans blue until 25 DPA (Fig. S1b-g), consistent with their role of assimilate transport to the developing endosperm. DNA fragmentation is uniformly detected in the mesocarp of *Brachypodium* pre-anthesis ovary (Fig. 1a). However, mesocarp cell disintegration proceeded most rapidly in lateral cells (Fig. 2a; Fig. S1c). Endosperm PCD indicated by positive Evans blue stain was detected by 15 DPA in a random pattern and increased in intensity thereafter (Fig. 1i,j; Fig. S1d). At 25 DPA, the entire endosperm was deeply stained except the aleurone layers. The progression of DNA fragmentation from pre-anthesis ovary to grain maturity is also shown by DNA laddering using gel electrophoresis (Fig. S2). There was an increase in the amount of less than 200 nt DNA fragments between from to pre-anthesis to 20 DPA. This suggests that DNA cleavage is happening throughout grain development though in different tissues. Less DNA fragments were detected at 25 DPA by which time the grain is mature and the tissues that remain viable; the embryo and aleurone, are not undergoing PCD.

#### Potential involvement of *MADS29* in *Brachypodium* PCD

Due to the importance of *OsMADS29* during rice grain PCD (Yin & Xue, 2012), we selected its ortholog in *Brachypodium* and analysed its localisation in developing *Brachypodium* grain using mRNA *in situ* hybridisation (Fig. 3). *BdMADS29* expression was detected in the endocarp and mesocarp cells above the ovule at pre-anthesis ovary (Fig. 3a-f). We found that *BdMADS29* generally localizes in cells undergoing PCD in developing *Brachypodium* grain and other tissues. Importantly, *BdMADS29* is also localized in *Brachypodium* endosperm cells undergoing PCD (Fig. 3g,h), in contrast, *OsMADS29* was not associated with rice endosperm PCD (Yin and Xue, 2012).

**Figure 3.**
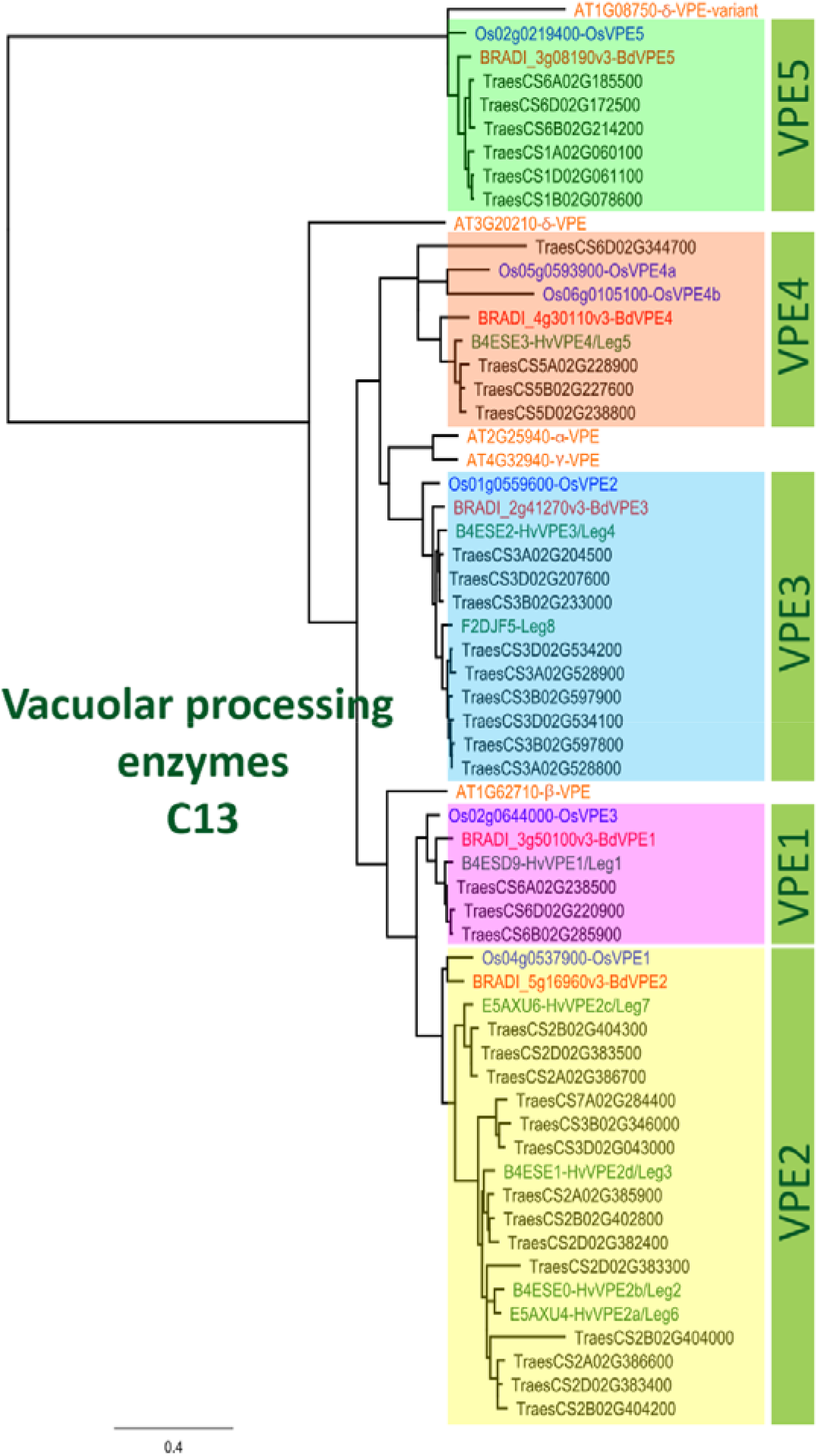
Localisation of BdMADS29 developing grain sections of *Brachypodium distachyon*. (a-d) Pr As Ov: pre-anthesis ovary, (e) mid-length grain at 6 DPA, (g) mature grain at 20 DPA, (a-c-e-g) antisense (b-d-f-h) sense. Arrows indicate the expression. An: anther, Fi: filament, NE: nucellar epidermis, Nu: nucellus paremchyma cells, Lo: lodicule, Mc: mesocarp, Pa: palea, Ec: endocarp, In: integument, En: endosperm, NL: nucellar lysate, Scale bar 100 μm.

### *Brachypodium* lacks expansion of VPEs found in *Triticeae*

Our sequence search identified 93, 48, 5, and 10 genes belonging to *Brachypodium* A1, C1, C13 and C14 protease family genes respectively (Table S1). Phylogenetic analyses incorporating protein sequences of these gene families from Arabidopsis, barley, rice and wheat enabled the identification and naming of putative orthologs in *Brachypodium*. Phylogenetic trees for A1, C1 and C14 families are presented in Figure S3, S4 and S5 respectively. Phylogenetic tree for C13 (Fig. 4) shows that monocot VPE protein sequences included in the analysis clusters into five clades with one *Brachypodium* VPE protein in each clade. Interestingly, we observed an expansion of *Triticeae* (represented by barley and wheat in the tree) sequences in VPE2, VPE3 and VPE5 clusters that is absent in *Brachypodium* and rice. Specifically, previous analysis by Radchuk *et al* 2010 leads us to suspect that the expansion of *Triticeae* protein sequences in VPE2 cluster may underlie differences in nucellus and nucellar epidermis PCD between barley and *Brachypodium*. Barley and wheat have by 4 and 14 sequences in the VPE2 clade respectively (Fig. 4). Three of the barley VPEs in that clade; *HvVPE2a, HvVPE2b* and *HvVPE2d* are most highly expressed at 4 days after flowering which coincides with nucellus PCD in barley endosperm fractions (Radchuk *et al* 2010). *BdVPE2* on the other hand clusters more closely to *HvVPE2c* whose expression was hardly detected in developing barley grains. We confirmed the expansion of VPE2 cluster by examining the ensembl Plants gene tree that contains *BdVPE2* (data not shown), which showed that the species in *Triticeae* tribe has more sequences in this clade than species in *Brachypodeae, Oryzinae*, and *Panicoideae*. In addition, *BdVPE2* (BRADI5g16960) expression in our transcriptome data was highest in mid-length and full-length grains (Fig. 5), a period that coincides with mesocarp and endosperm PCD and not nucellus, therefore suggesting a functional difference with barley HvVPE2 genes.

**Figure 4.**
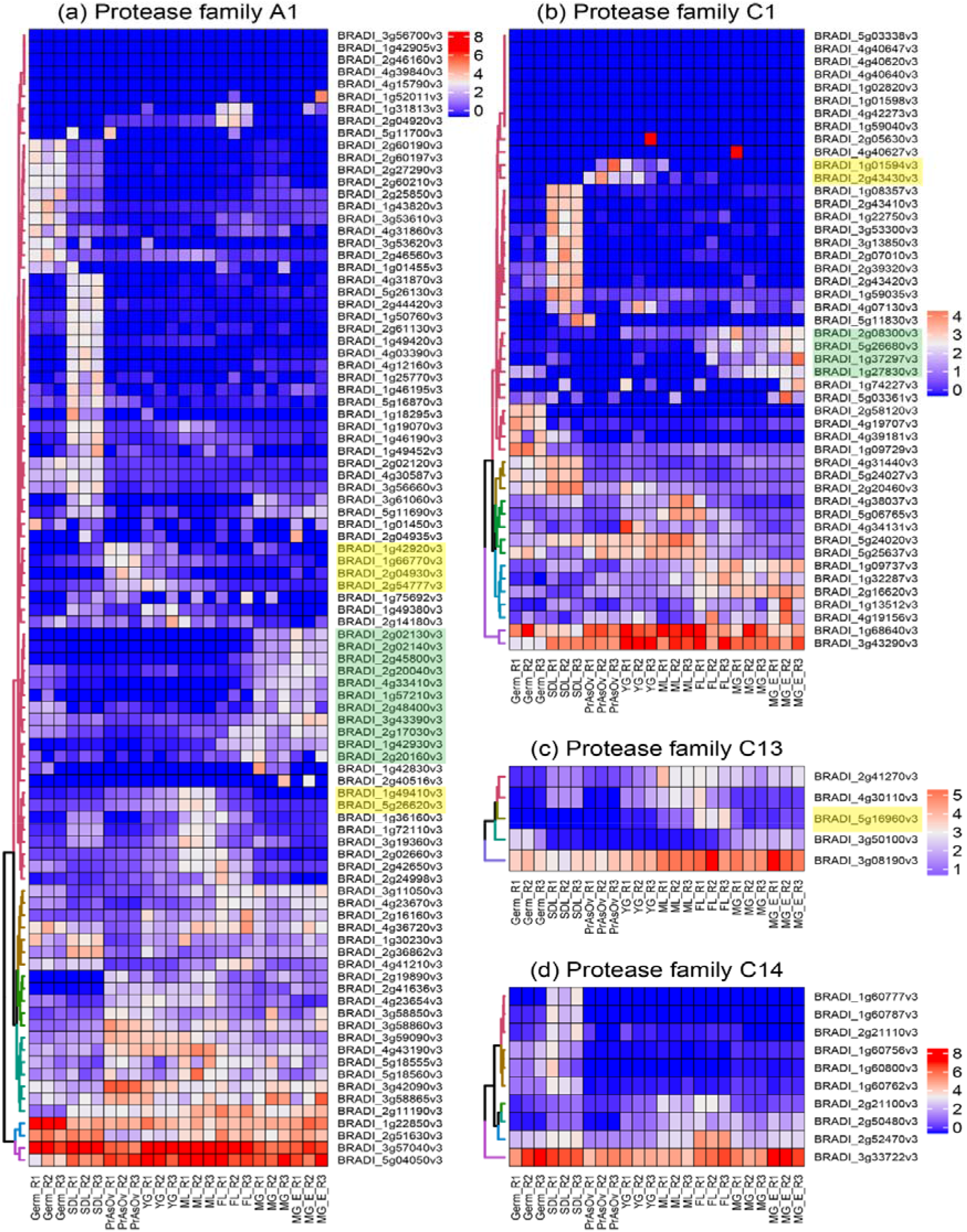
Phylogenetic Tree of Vacuolar Processing Enzymes C13 family. It illustrate the five groups VPE1-5 and includes VPEs proteins from *Arabidopsis thaliana, Oryza sativa, Brachypodium distachyon, Hordeum vulgare* and *Triticum aestivum*. JTT+I was used as protein model and the sequences were aligned with ClustalW, then inferred using Maximum Likelihood method with a bootstrap of 1,000 replicates. The tree was generated using MEGA10.

**Figure 5.**
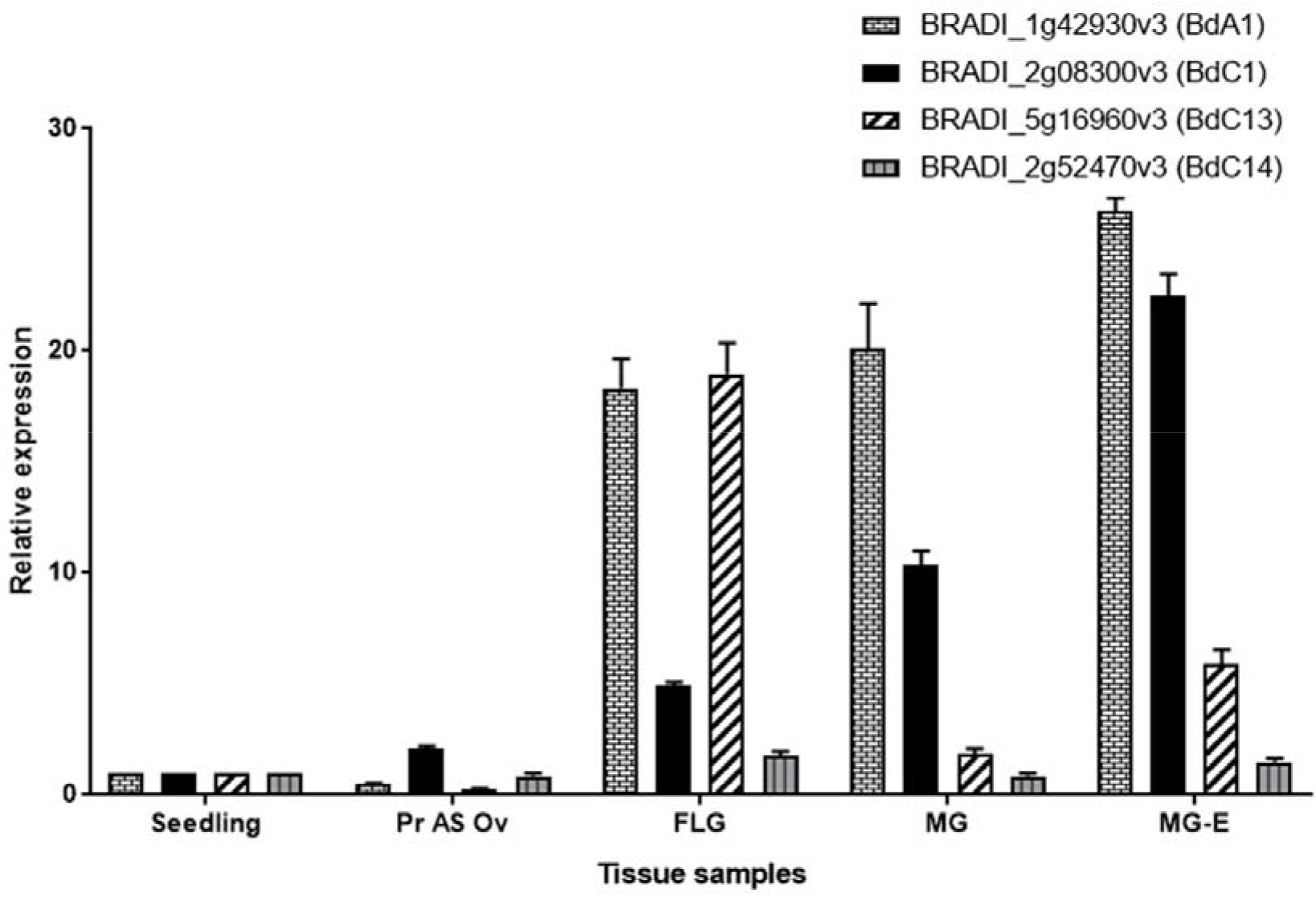
Hierarchically clustered expression profile of protease genes from A1, C1, C13 and C14 families in vegetative and grain tissues of *Brachypodium*. Heatmap was plotted with DESeq2 normalized RNA-Seq reads centred with unit variance scaling. Genes in clusters highlighted are active in nucellar and mesocarp (yellow), and endosperm (green). PrAnOv, pre-anthesis ovaries; YG, young grain (1-3 DPA); ML, mid-length grain (3-8 DPA); FL, full-length grain (8-15 DPA); MG, mature grain (15-20 DPA); MGE, mature grain without embryo; Germ, germinating grain; SDL, seedling 3-4 days after germination.

### Expression analysis identifies putative grain specific proteases

Figure 5 shows DESeq2 normalized, autoscaled RNA-Seq expression profile of protease genes belonging to A1, C1, C13 and C14 families. The samples can be broadly divided into vegetative tissues (seedling) and grain tissues (germinating grain, pre-anthesis ovary, young grain, mid-length grain, full-length grain and mature grain). The hierarchically clustered expression profile revealed clusters of A1, C1 and C13 proteases that are specifically expressed in developing *Brachypodium* grain tissues. Although, proteases belonging to the C14 family are expressed in grain tissues, they were also highly expressed in vegetative tissue and none was judged to be grain specific. However, clusters of proteases that are highly expressed during the period of nucellus and mesocarp PCD were identified in A1, C1 and C13 families (highlighted yellow in Fig. 5a,b,c). Also clusters of genes that are highly expressed in the endosperm can distinguished in A1 and C1 (highlighted green in Fig. 5a,b). We selected one representative gene from each protease family and validated their expression using quantitative RT-PCR (Fig. 6). The results agree with the RNA-Seq expression profile.

**Figure 6.** Normalised relative RT-qPCR expression of selected peptidase candidates, in seedling, pre-anthesis ovary (Pr As Ov), full-length grain (FL), mature grain (MG) and mature grain without embryo (MG-E). Seedling was used as calibrator. Error bars indicate ± SD (n = 3). ANOVA showed significant expression difference in between samples for all genes tested at P < 0.05.

## Discussion

Plant PCD research in the past three decades have demonstrated the importance of timely death of certain cells for the normal development of plants. Current knowledge of PCD in developing grass grain are mainly derived from studies of domesticated species. Disruption of normal grain developmental PCD have been shown to adversely affect grain filling in barley, maize and rice (Young *et al*., 1997; Yin & Xue, 2012; Radchuk *et al*., 2018).

In this study, we investigated grain PCD in a wild species, *Brachypodium*. While the general features of PCD are similar between developing cereals and *Brachypodium* grains, we also observed subtle differences in timing, pattern and progression of PCD between cereals and *Brachypodium* grains. For example, we detected PCD in pre-anthesis ovary of *Brachypodium* (Fig. 1a). PCD in pre-anthesis ovary have not been previously reported in other species. A possible reason is that most studies focus on PCD post fertilization. Our results suggest that PCD is activated in *Brachypodium* nucellar and mesocarp cells before fertilization. Radchuk *et al*, (2010), detected PCD in barley nucellus and mesocarp at anthesis (0 DPA) and 6 DPA respectively. Their results does not exclude the possibility of nucellar PCD already taking place before the sampled time point. However, it does suggest that barley mesocarp PCD is initiated between 2 and 6 DPA. Compared to *Brachypodium* therefore, it appears that initiation of mesocarp PCD occurs later in barley compared to *Brachypodium*.

*Brachypodium* nucellar disintegration occurs in a centrifugal pattern similar to barley, rice and wheat. However, the rate of nucellar cells disintegration is slow compared to barley. Rapid disintegration of the nucellus is thought to provide growth space and resources for the incipient endosperm in cereals (Domínguez & Cejudo, 2014). In maize, early endosperm development around the period of nucellar PCD influences final grain size (Leroux *et al*., 2014). It is therefore possible that slow nucellar disintegration in *Brachypodium* impedes its early endosperm development because of reduced space and poor resource supply from degenerated cells. Moreover, *Brachypodium* nucellar epidermis enlarges greatly before undergoing PCD ca. 6 DPA. Interestingly, the nucellar epidermis does not collapse or disintegrate throughout grain development. We have previously shown that the nucellar epidermis serves as assimilate transport channel towards the endosperm (Solomon & Drea, 2019). A similar role is played by rice nucellar epidermis (Oparka & Gates, 1981), which in contrast to *Brachypodium* nucellar epidermis collapses during grain development.

Conversely, mesocarp disintegration occurs at a faster rate in *Brachypodium* compared to barley. This agrees with the earlier observation that mesocarp PCD is initiated earlier in *Brachypodium* compared to barley. Because barley mesocarp PCD is only detected after endosperm cellularisation, Radchuk *et al* (2010), proposed that barley mesocarp PCD is coordinated with endosperm development. In maize, PCD in nucellar placento-chalazal region is coordinated with endosperm cellularisation and is completed before the commencement of major grain filling (Kladnik *et al*., 2004). Okada *et al*, 2017, confirmed coordination between endosperm development and mesocarp PCD in wheat and barley. They showed that mesocarp cells of unfertilized ovary do not disintegrate. Instead mesocarp cells swell laterally and force the lemma and palea apart in an apparent attempt to increase the chances of cross pollination and fertilization. Our results suggests that *Brachypodium* lacks this coordination of endosperm development with mesocarp PCD. Furthermore, the rate of degeneration of *Brachypodium* grain mesocarp cells is most rapid at the lateral regions (Fig. 2a). This contrasts with the centrifugal mesocarp degeneration pattern reported in barley, rice and wheat (Domínguez and Cejudo, 2014; Dominguez *et al*., 2001; Radchuk *et al*., 2010; Yin and Xue, 2012). Because the endosperm subsequently expands laterally to fill the space left by disintegrated mesocarp cells, we speculate that the pattern of mesocarp disintegration may contribute to the flat shape of *Brachypodium* grains.

*Brachypodium* endosperm PCD was detected at 15 DPA (Fig. S1g) and did not progress in any discernible pattern. Random progression of endosperm PCD has also been observed in wheat, whereas the maize endosperm PCD proceeds in an organized top-to-base fashion. The difference in the pattern of endosperm PCD has been attributed to grain size. It has been suggested that while PCD can proceed randomly in the comparatively smaller endosperm of wheat, larger of endosperm of maize require PCD to be executed in organized manner (Young and Gallie, 2000, 1999; Young *et al*., 1997).

Proteases are known to contribute to the execution of PCD. Although the details of the contribution of individual genes are yet to be elucidated, the expression profile of several protease genes strongly coincide with PCD in different grain tissues (Buono *et al*., 2019). Such correlative evidence have been used to implicate a number of protease genes as actors during PCD of one or more cereal grain tissues. To gain a comprehensive view, we identified all *Brachypodium* genes that belongs to protease families whose member have been implicated in PCD of developing grains. The families comprise A1, C1, and C13. No reports exist yet that links family C14 (metacaspases) to grain PCD. However, we included family C14 in our analyses because members of the family have been characterised in detail in *Arabidopsis* (Tsiatsiani *et al*., 2011). Furthermore, the expression of metacaspase genes were reduced in *Brachypodium* callus treated with 5 and 50 μM 5-azacitidine (Betekhtin *et al*., 2018). Phylogenetic analyses enabled the identification of *Brachypodium* orthologs of the selected families. An RNA-Seq survey of the expression profile of genes in the selected families revealed that majority of genes in A1 and C1 are lowly expressed in the vegetative and grain tissues sampled. The analyses also revealed novel candidate genes that may be further explored for their roles in grain PCD (Table 1). Nevertheless, mRNA expression of proteases may not always mean activity because proteases are known to be inhibited by cystatins (Subburaj *et al*., 2017).

**Table 1.** Candidate protease genes for *Brachypodium* grain PCD

| Family | Family | Nucellar | Mesocarp | Endosperm |
| --- | --- | --- | --- | --- |
| A1 | BRADI1g42930 | - | - | YES |
| A1 | BRADI2g45800 | - | - | YES |
| A1 | BRADI1g49410 | YES | YES | - |
| A1 | BRADI5g26620 | YES | YES | - |
| A1 | BRADI2g02140 | - | - | YES |
| C1 | BRADI2g08300 | - | YES | YES |
| C13 | BRADI5g16960 (BdVPE2) | - | YES | YES |

The protease family C13 (VPEs) remarkably has genes that are the most highly expressed in the four families surveyed. Detailed expression analysis of VPEs in barley grain fractions revealed identical expression profile of *HvVPE2a, HvVPE2b* and *HvVPE2d* in nucellar and endosperm fractions (Radchuk *et al*., 2010). *HvVPE2a* (nucellain) localizes in degenerating nucellar cells (Linnestad *et al*., 1998). Therefore, these three genes are hypothesized to play major roles in barley grain nucellar PCD. The three genes and their wheat orthologs, form a distinct subclade (Fig. 4). However, *BdVPE2* is more similar to *HvVPE2c*, which is barely expressed in barley grains. We speculate that the combined activities of barley *HvVPE2a, HvVPE2b* and *HvVPE2d* facilitates the rapid disintegration of the barley nucellus. *Brachypodium* on the other hand lacks direct ortholog of these genes and shows slow nucellar degeneration rate. In addition, *BdVPE2* expression does not overlap nucellar PCD. This prompts us to suggest that the slow progression of *Brachypodium* nucellar PCD and the persistence of the nucellar epidermis may be partly due to lack of VPEs that facilitate nucellus disintegration.

## Conclusion

Our study provides details of grain PCD in a wild species, *Brachypodium*. The results and discussion, presented in a comparative perspective to what is already known of cereal grain PCD highlights similarities and crucially, differences in the timing, pattern and progression of grain PCD between *Brachypodium* and cereals. It appears that the rapid degeneration of the nucellar after fertilization in cereals stimulates rapid endosperm expansion, resulting in large grains. We suspect that reduced expansion stimulus offered by slow degenerating *Brachypodium* nucellus contribute to its small grain size. Our suspicion is supported by *Brachypodium* having just one (*BdVPE2*) ortholog of nucellus degenerating VPE, whereas barley and wheat has four and fourteen respectively. The lateral pattern of *Brachypodium* mesocarp degeneration appears to contribute to its dorsi-ventrally flattened grain shape.

## Acknowledgements

This research was initiated as part of BBSRC grant BBI01215X/1 previously held by SD CUS is supported by TETFUND, Nigeria.

## Author Contribution

CUS and SS designed the study and performed all experiments and analyses under the supervision of SD. CUS and SS wrote the manuscript. All authors read and approved the final manuscript.

